# Generic transcriptional response of *E. coli* to stress

**DOI:** 10.1101/020578

**Authors:** Ranginee Choudhury, Saroj Kant Mohapatra

## Abstract

Bacteria are often exposed to various stressors with inter-linked effects leading to cross-resistance. In order to study the common molecular response to stress in bacteria, we collated and compared transcriptional response of *E. col*i under a variety of biotic and abiotic stresses. Bacterial genome-wide gene expression data were retrieved from the NCBI Gene Expression Omnibus (GEO) database and interrogated to identify differentially expressed genes.

One hundred and sixty eight genes displayed distinct transcriptional response of *E. coli* to stress, with simultaneous down-regulation of flagellar assembly pathway and up-regulation of the global regulator rpoS.

This computational analysis summarizes stress responsive genes in *E. coli* and their inter-relationships.

---

Antibiotic resistance in a global problem of high magnitude and demands a comprehensive approach. Antimicrobials activate stress responses in bacteria which contribute to development of resistance^[1]^. Response to a particular stressor may promote resistance to other unrelated stresses. This involves metabolic rewiring of the organisms exposed to stress^[2]^. The metabolic changes are often transient but occasionally permanent depending on the adaptive changes in the genetic architecture. It is, therefore, pertinent to study the stress responses which are thought to be resistance determinants^[1]^.

We conducted computational analysis of genome-wide transcriptional response of *E. coli* to stress. Systematic search of National Center of Biotechnology Information (NCBI) Gene Expression Omnibus (GEO) was conducted to select studies on transcriptional response of *E. coli* to antibiotic and other stresses. The selected stressors represented different groups of antibiotics, i.e., inhibitors of protein synthesis (e.g., tetracycline), cell wall (e.g., cefsulodin), nucleic acid (e.g., nalidixic acid) and folate (e.g., sulfisoxazole); biocides, such as, copper, benzalkonium hydrochloride (BC), polyhexamethylene biguanide (PHMB), hydrochloric acid (HCl) and ethanol. Food additives such as common salt, glycerol, acetic acid and other stressors, such as, temperature and ethidium bromide were also included. PHMB and BC were selected in view of their extensive use as disinfectants in the health care facilities, considered hot-spots of emergence and transmission of antibiotic resistance genes^[3]^.

Biocide stress is of importance because there are several reports of cross-resistance between quaternary ammonium compounds and antibiotics primarily mediated by polyspecific multidrug pumps^[4]^. Two studies on gene-expression induced by BC were included. In one of the studies the dosage was sub-inhibitory (7 and 9 *μ*g/ml) and in the other *E. coli* was progressively adapted to 50 *μ*g*/*ml, a concentration much higher than its minimum inhibitory concentration (12 *μ*g/ml)^[5]^. Concentration of PHMB used in the selected study was within the range of bacteriostatic concentration which is reported to be 1-10 *μ*g/ml ^[6]^, while copper concentration was near lethal at 2mM ^[7]^. The bactericidal concentration of ethanol is 60%-90% solution in water (volume/volume) ^[8]^. In the selected study 5% ethanol was used as a stressor. No data were available for the concentration of antibiotics from one of the studies (GSE10440).

Datasets from six studies (GSE1780, GSE2827, GSE6712, GSE10159, GSE10440 and GSE11041, available at http://www.ncbi.nlm.nih.gov/geo/) were selected (Table 1). Each dataset (except GSE10440) was accompanied by a research publication. In all, the six studies included 32 stressors (Table 2).

**Table 1:**
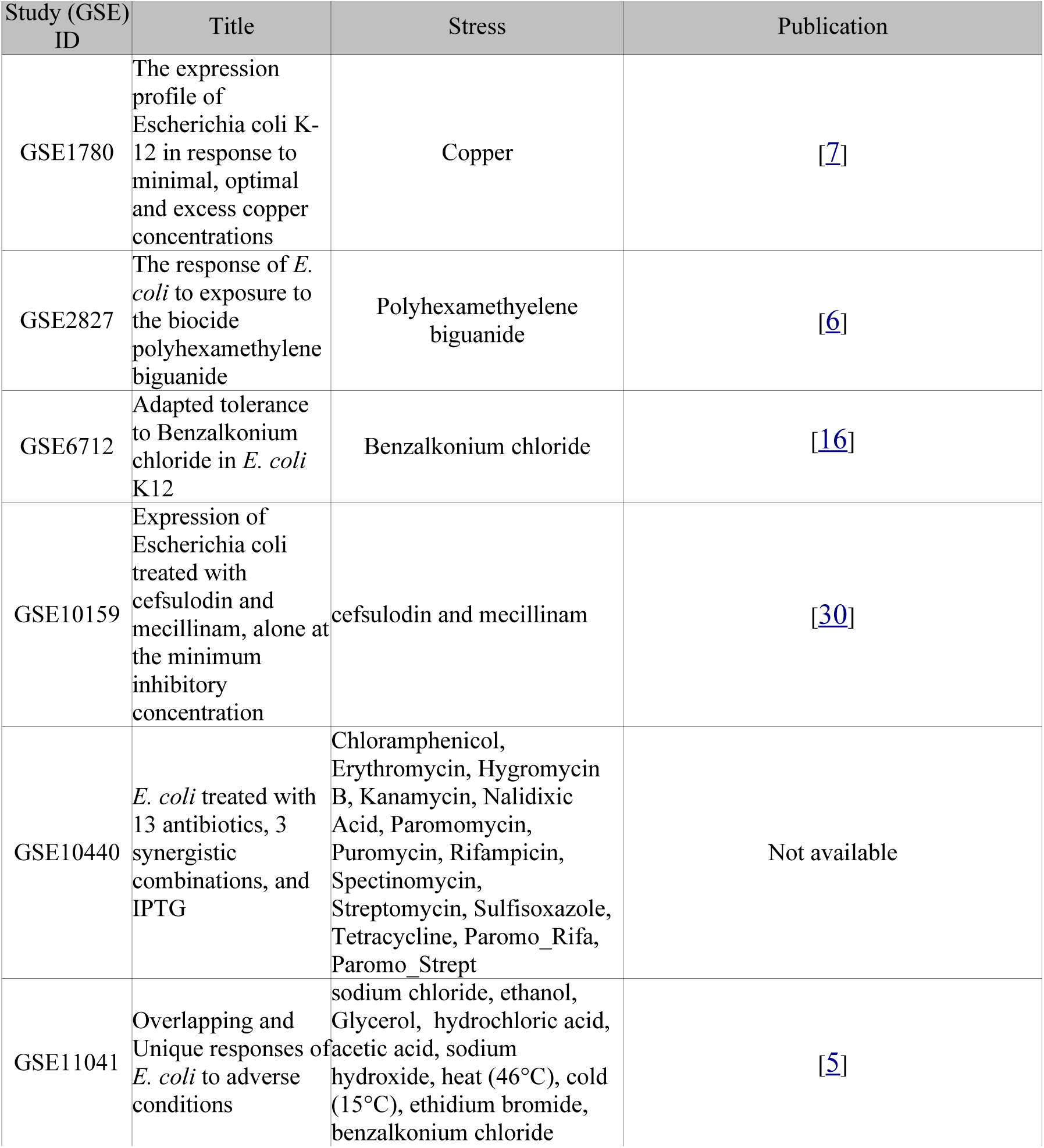
Characteristics of the six data sets included in the current study

**Table 2.** List of stressors

| Study ID | Stress | Description |
| --- | --- | --- |
| GSE1780 | Copper | E coli K12 response to high copper concentration (2mM) compared with no added copper to the growth medium. |
| GSE10159 | cefsulodin60_5min | E coli K12 strain MG1655 cells treated with cefsulodin concentration of 60µg/ml for 5minutes and compared with untreated cells. |
| GSE10159 | cefsulodin60_10min | E coli K12 strain MG1655 cells treated with cefsulodin concentration of 60µg/ml for 10 minutes and compared with untreated cells. |
| GSE10159 | mecillinam0.3_60min | E coli (MG1655) cells treated with mecillinam concentration of 0.3mg/ml for 60 minutes and compared with untreated cells. |
| GSE10440 | Chloramphenicol | Not available |
| GSE10440 | Hygromycin B | Not available |
| GSE10440 | Nalidixic Acid | Not available |
| GSE10440 | Puromycin | Not available |
| GSE10440 | Spectinomycin | Not available |
| GSE10440 | Sulfisoxazole | Not available |
| GSE10440 | Paromo_Rifa | Not available |
| GSE10440 | Erythromycin | Not available |
| GSE10440 | Rifampicin | Not available |
| GSE10440 | Streptomycin | Not available |
| GSE10440 | Tetracycline | Not available |
| GSE10440 | Paromo_Strept | Not available |
| GSE10440 | Kanamycin | Not available |
| GSE10440 | Paromomycin | Not available |
| GSE11041 | NaCl | *E. coli* K12 MG1655 response to 4.5% sodium chloride compared with control. |
| GSE11041 | HCl | *E. coli* K12 MG1655 response to hydrochloric acid (pH 4.75 ± 0.05) compared with control. |
| GSE11041 | CH3COOH | *E. coli* K12 MG1655 response to acetic acid (pH 5.9 ± 0.05) compared with control. |
| GSE11041 | EtOH | *E. coli* K12 MG1655 response to 5% ethanol compared with control. |
| GSE11041 | BC7 | *E. coli* K12 MG1655 response to 7 µg/ml benzalkonium hydrochloride compared with control. |
| GSE11041 | EtBr | *E. coli* K12 MG1655response to 150 µg/ml ethidium bromide compared with control. |
| GSE11041 | Glycerol | *E. coli* K12 MG1655response to 15% glycerol compared with control. |
| GSE11041 | NaOH | *E. coli* K12 MG1655response to acetic acid (pH 9.6 ± 0.05) compared with control. |
| GSE11041 | deg46 | *E. coli* K12 MG1655 response to high temperature (46 °C) compared with control. |
| GSE11041 | deg15 | *E. coli* K12 MG1655 response to low temperature (15 °C) compared with control. |
| GSE11041 | BC9 | *E. coli* K12 MG1655 response to 9 µg/ml benzalkonium hydrochloride compared with control. |
| GSE2827 | PHMB | *E. coli* K12 MG1655 response to 7.5 mg/l benzalkonium hydrochloride compared with control. |
| GSE6712 | BC | Transcriptomic response of *E. coli* K12 gradually adapted to benzalkonium (BC) 7-8 times the initial MIC. The bacteria were grown in the absence of BC. |
| GSE6712 | BC50 | Transcriptomic response of *E. coli* K12 gradually adapted to benzalkonium (BC) 7-8 times the initial MIC. The bacteria were grown in the presence of BC (50 µg/ml). |
| Table 2: List of stressors |  |  |

The data sets were generated using five different genome-wide gene expression platforms (GPL1246, GPL3154, GPL6679, GPL534 and GPL189), with different number of probes per array (ranging from 4290 to 14400, Figure 1). This discrepancy in number of probes per study was resolved by mapping the probes to the *E. coli* K12 genome (3110 genes) and conducting subsequent analysis at the gene-level. For each group, relative gene expression of bacteria under stress was calculated with respect to the control state (i.e., no stress). In all, 32 comparisons were performed, one for each of the stresses listed in Table 2 Thus for each gene, 32 relative expression values were computed. We reasoned that for a gene to be part of the generic stress response, it should behave similarly across multiple stresses, i.e., it should be either mostly up-regulated or mostly down-regulated for various stress conditions. This was tested by conducting one-sample t-test for deviation of relative gene expression from zero (where zero represents no change in gene expression in the logarithmic scale). Correction for multiple testing was performed using the method of Benjamini and Hochberg. Any gene associated with p-value of less than 0.05 was included in the list of 168 common genes differentially expressed in response to stress, representing the generic response to stress (Table 3, Table S1).

**Figure 1.**
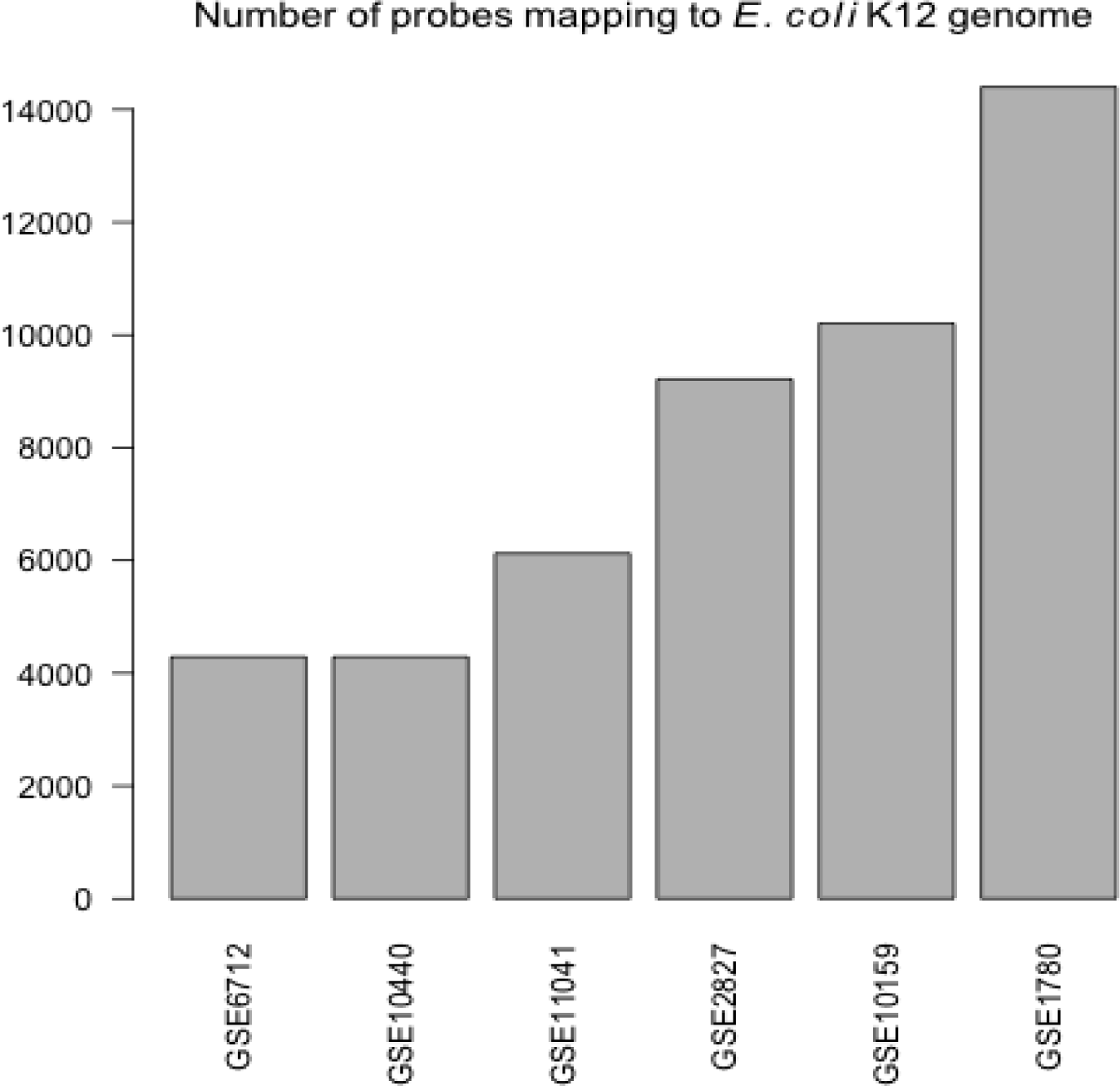
The number of probes in each study that are mapped to *E. coli* K12 genome. Both GSE6712 and GSE10440 map to the same platform ID GPL189 (Sigma Genosys Panorama Escherichia coli Gene Array) and report the same number of probes (4290).

**Table 3.**
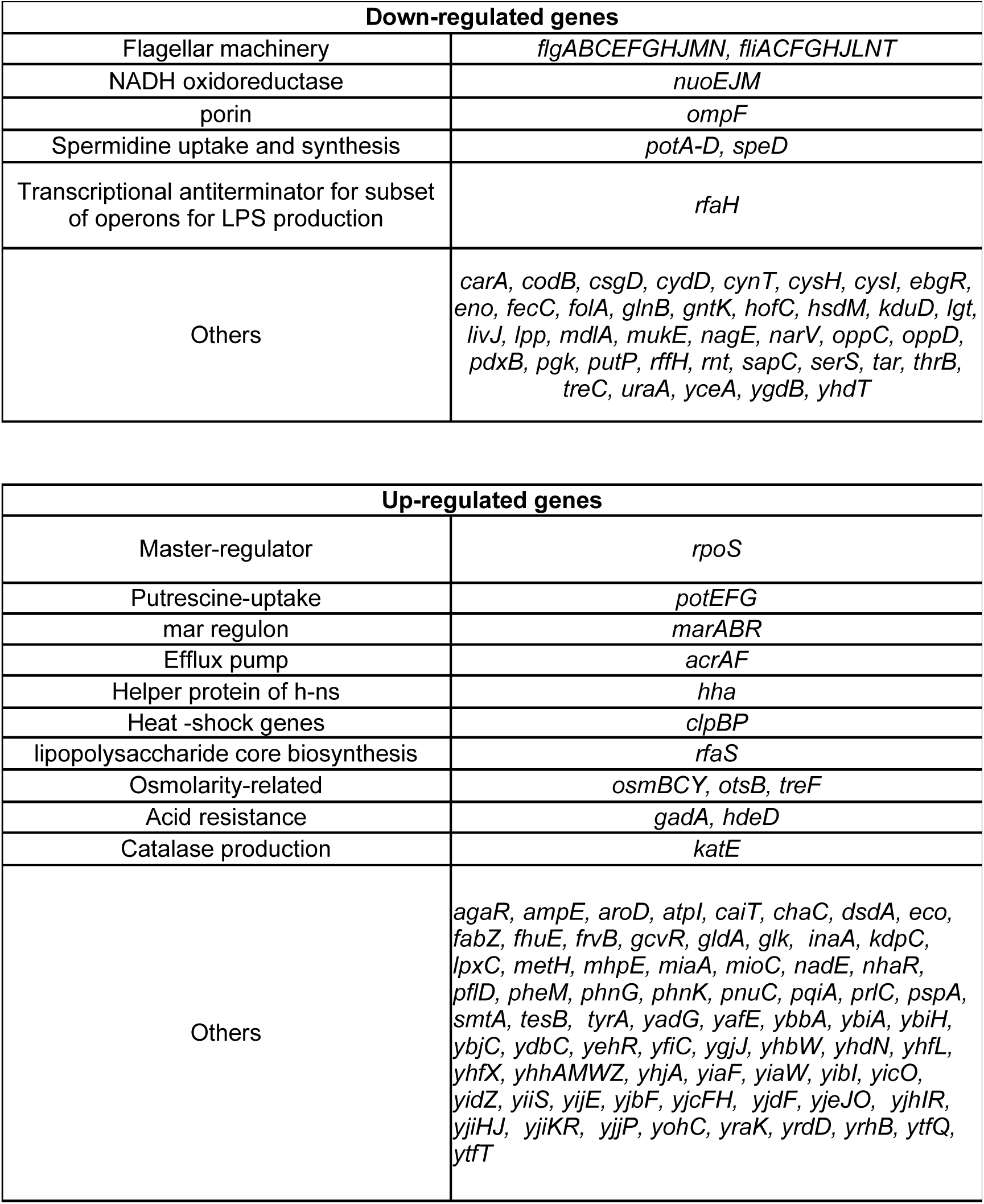
List of up and down-regulated genes.

The list of 168 genes included both up-regulated (rpoS, mar regulon, hha, clpB, pspA, osm group, gadA, katE, acrA, acrF, and the putrescine transporters potE-G, etc.) and down-regulated genes (including flg and fli groups, ompF, nuo group, and the genes for spermidine uptake potA-D). Up-regulation of marA was observed under aminoglycoside, sulfisoxazole, ethidium bromide, heat, alcohol, acid and glycerol stresses. BC-adapted strains showed up-regulation of marA, possibly related to the mar-sox-rob network, although the other two genes did not reach the level of significance as marA. Up-regulation of genes involved in salt stress was observed. Osmotically inducible genes – osmY, osmB, osmC and otsB ^[9]^ were over-expressed under the influence of many stressors.

Characteristic clusters of both genes and stress conditions were observed on the heat map of the 168 differentially expressed genes (Figure 2). Two clades of affected genes showed down-regulation of flagellar biosynthesis pathway, NADH-ubiquinone oxido-reductase, porin, spermidine synthesis/uptake and, simultaneous up-regulation of rpoS, putrescine uptake, mar regulon and efflux pumps. Four distinct clades of stressors were obtained. These can be broadly grouped as antibiotics clustering with excess copper forming two sub-clusters and biocides, temperature, salt, acid stress grouping together. PHMB formed a cluster by itself.

**Figure 2.**
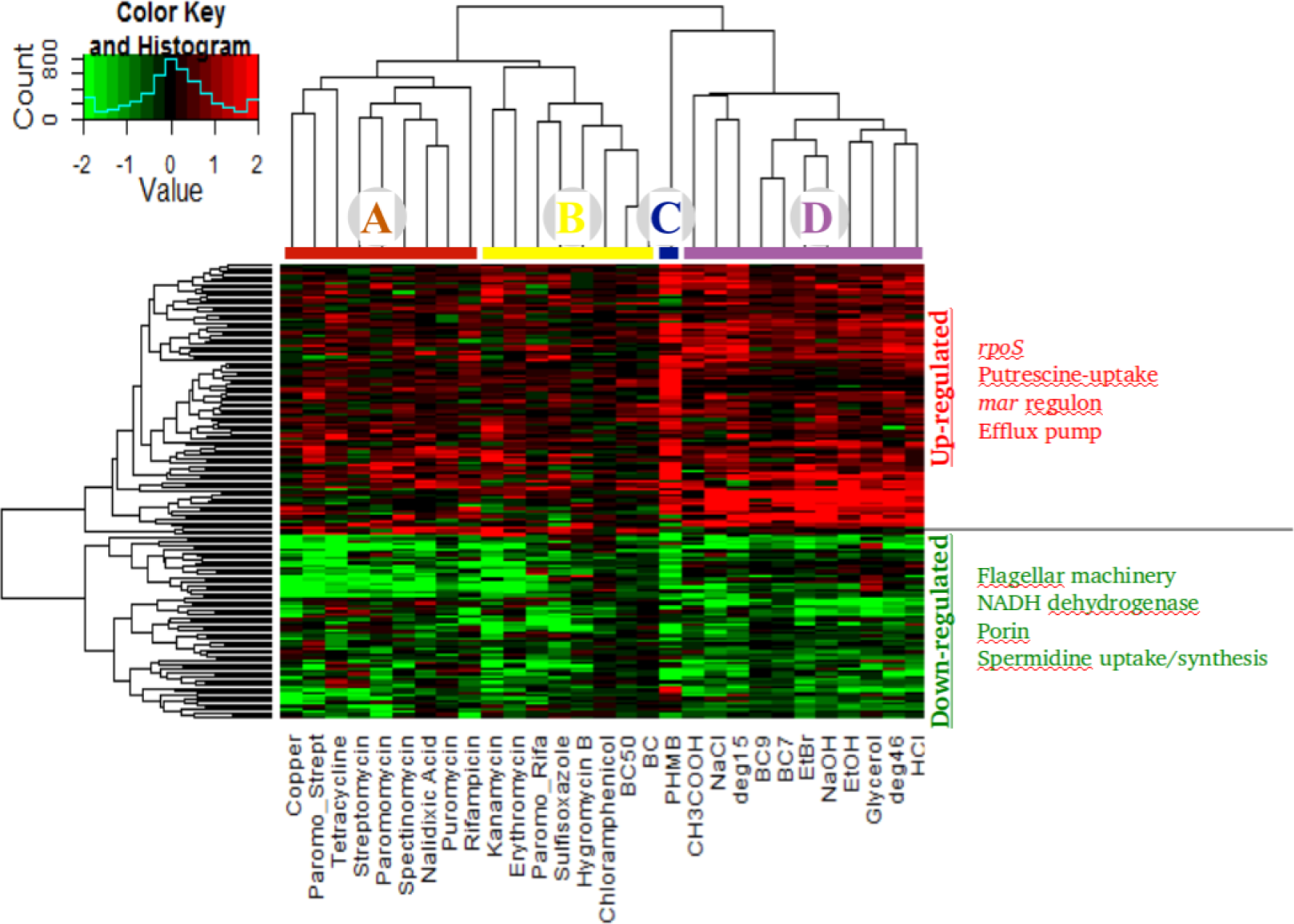
Heatmap of 168 genes which are differentially regulated in response to the spectrum of stressors. Up-regulated genes include the master regulator rpoS, genes for putrescine uptake, mar regulon, and the efflux pump genes. Down-regulated genes include porin, NADH ubiquinone oxido-reductase 1, genes for flagellar synthesis, spermidine uptake and synthesis. Excess copper co-clusters with antibiotics, while PHMB forms a subcluster distinct from other biocides.

Our analysis revealed that excess copper, which is used as a biocide, elicits transcriptional stress response that is similar to antibiotics, especially, protein synthesis inhibitors (PSIs) (Figure 2). This is interesting since their mechanisms of action differ. Copper toxicity leads to perturbation in protein function owing to its competition with other metal binding sites in the proteins ^[7]^. PSIs mainly act on the ribosomes and jeopardise the protein synthesis machinery. However, both the stresses induce Fenton-mediated hydroxyl radical formation that utilize internal iron of the Fe-S clusters ^[7, 10]^. In our study, down-regulation of NADH-ubiquinone oxidoreductase (nuo) pathway is observed as a result of the excess copper and PSIs (Table S1). Kohanski and colleagues (2007) ^[10]^ observed up-regulation of the NADH-ubiquinone oxidoreductase (nuo) pathway in response to antibiotic stress and attributed it to generation of hydroxyl ions resulting in the killing of bacterial cells. However, nuo genes were reported to be repressed under ampicillin and ofloxacin stress ^[11]^. Down-regulation of nuo genes is a protective mechanism adopted by the *E. coli* cells to prevent burst of superoxide generation via the respiratory chain ^[9]^.

We sought to unearth important transcriptional patterns from a combined gene expression dataset. These patterns suggested some mechanisms that may prepare the organism to confront current stress and surpass future challenges. Our salient findings follow.

## 1. Flagellar assembly and rpoS

Enrichment analysis of the 168 genes revealed flagellar assembly as the most repressed pathway in response to stress (Figure S1). Expression of the genes encoding flagellar proteins and motor components are highly regulated ^[12]^, with participation of several operons ^[13]^. These genes are negatively controlled by RpoS, which, in turn, acts either through the regulation of the flagellar master regulator FlhCD, or the flagellar sigma factor FliA ^[14]^. We observed up-regulation of rpoS in response to a number of stresses, namely, protein synthesis inhibiting antibiotics (PSIs), sulfisoxazole, salt, hydrochloric acid, sodium hydroxide, acetic acid, heat and cold stress, ethanol, BC7 and BC9; and simultaneous down-regulation of flagellar assembly gene expression (Figure 3) – consistent with the negative control exerted by RpoS. The coordinated down-regulation of this pathway is perhaps an adaptive response that provides survival advantage to bacteria under stress by restriction of movement and conservation of energy.

**Figure 3.**
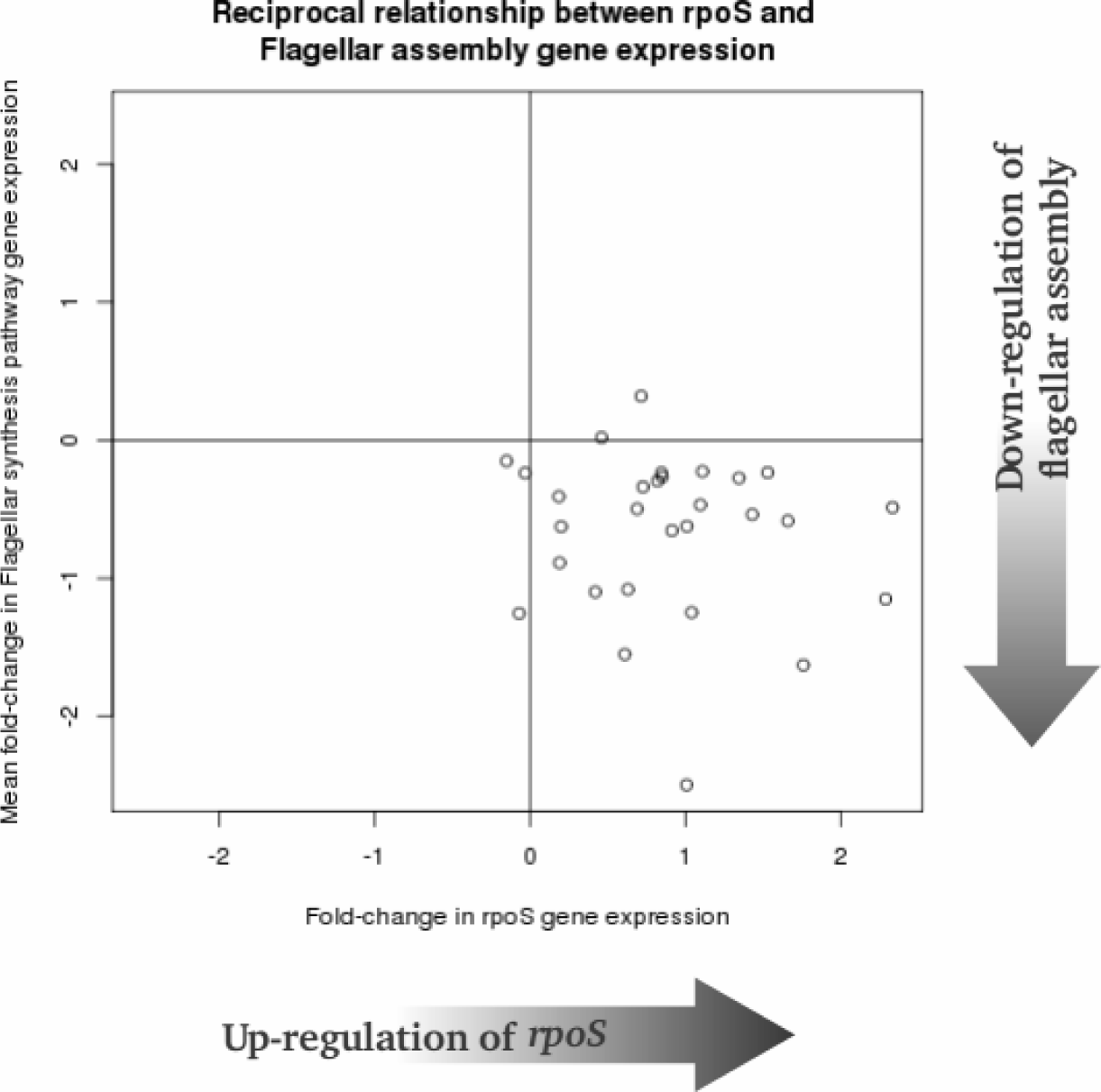
This scatter-plot shows reciprocal relationship between expression of rpoS and genes of Flagellar assembly. Log-fold changes in gene expression are plotted: x-axis for rpoS under stress, y-axis corresponds to average log-fold change in gene expression of flagellar assembly pathway under stress. Each point corresponds to a particular stress. The plot area is divided into four quadrants. Clustering of all points in lower right suggests simultaneous up-regulation of rpoS and down-regulation of Flagellar assembly, consistent with the regulatory role of rpoS.

## 2. Influx and efflux

Antibiotic resistance, which appears with the activation of mar regulon, is accomplished by preventing influx of toxic substances into the cells and also by enhancing efflux of the same. The former is accomplished partially by repressing ompF and the latter is achieved by over-expressing acr-tolC ^[15]^. We observed repression of ompF in response to a number of stresses (excess copper, nalidixic acid, paromomycin, puromycin, rifampicin, streptomycin, tetracycline, PHMB, salt, glycerol, hydrochloric acid, ethanol, BC7 and BC); and up-regulation of acrA in response to kanamycin, paromomycin, sulfisoxazole, tetracycline, and benzalkonium hydrochloride adaptation (Table 3; Table S1). Up-regulation of acrAB-tolC in BC adapted cells, likely caused activated efflux and contributed to the survival and adaptation of *E. coli* to high BC. This also explains resistance to multiple antibiotics shown by the adapted strain ^[16]^. It is worth noting here that mar locus has been proposed as a “stepping stone” to progressively increasing resistance, facilitated by its role in efflux thereby contributing to survival advantage, that in turn result from subsequent adaptive mutations elsewhere on the chromosome ^[15]^.

## 3. Polyamine uptake

Previous studies have shown the role of polyamines in reducing oxidative stress in *E. coli*^[17]^. Spermidine and putrescine are pre-dominant cytoplasmic polyamines in bacteria ^[18]^. We observed over-expression of genes responsible for putrescine uptake – potEFG; and repression of genes responsible for spermidine uptake – potABCD ^[19]^ (Table 3, Figure 4). Additionally repression of speD gene encoding S-adenosyl methionine (SAM) decarboxylase was observed. This enzyme converts SAM into decarboxylated SAM, which acts as a cofactor for spermidine biosynthesis ^[19]^. It is possible that *E. coli* responds to stress by limiting intracellular spermidine concentration and increasing putrescine concentration. At higher concentration spermidine is reported to arrest protein synthesis during stationary phase of *E. coli*^[20]^. Moreover, spermidine carries 3+ charge, and plays a role in stabilization of the outer membrane against antimicrobial attack. In contrast, intracellular putrescine or cadaverine carry a charge of 2+ each ^[18]^, and protect the bacterial cell from oxidative stress by stimulating the oxyR expression ^[21]^. This probably explains the need for maintaining an exogenous pool of spermidine and endogenous pool of putrescine in bacteria under stress, and appropriately regulated at the level of transcription, as shown. Furthermore, putrescine has been earmarked as an important communicating molecule in drug resistance ^[22]^. Experimental testing of this finding by generating specific mutants is beyond the scope of this study. However, this finding is consistent with earlier findings in terms of spermidine toxicity ^[20, 23]^ and does open up possibility for further explorations on this aspect.

**Figure 4.**
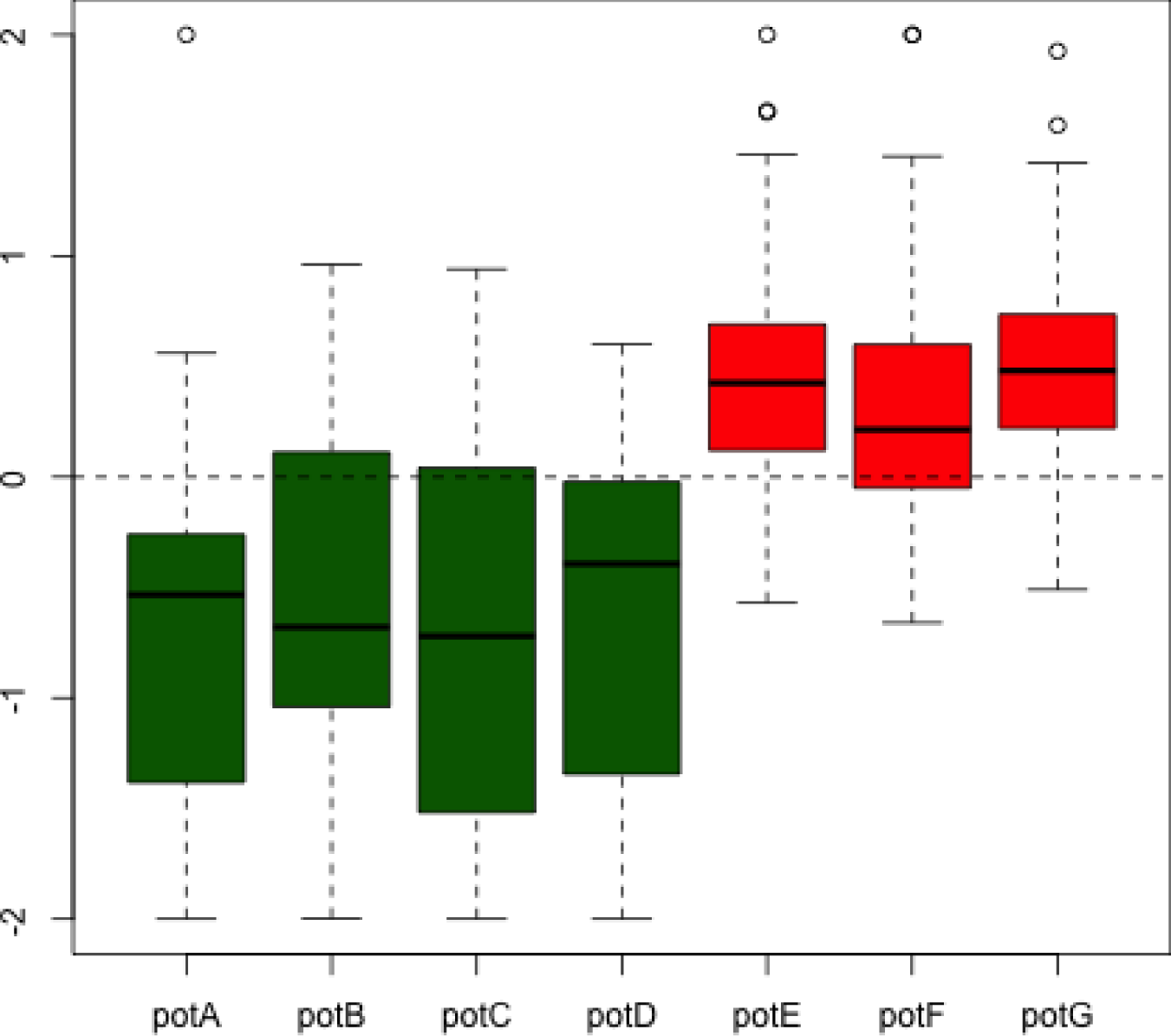
Box plots of relative gen expression of polyamine transporters in response to stressors. Genes for spermidine uptake – potABCD are down-regulated (shown in green), whereas genes for putrescine uptake – potEFG are up-regulated (shown in red).

Our results provide relevant insights about generic response of *E. coli* to a variety of stresses. The compelling theme appears to be metabolic control mediated by RpoS and regulation of flagellar machinery. Energy expensive flagellar machinary is functionally linked to NADH-ubiquinone oxido-reductase (ndh-1) of the electron transport chain. Oxidation of NADH is converted into electrochemical proton gradient by ndh-1 which plays an important role in flagellar motility in *E. coli*^[24, 25]^. Our results show down-regulation of both of these pathways establishing bacterial adaptation to stress via energy-conserving pathways. Modulation of intracellular concentration of polyamines might be an important factor for adaptation to stress. Also, this is consistent with the earlier reports on polyamines playing a crucial role in the regulation of rpoS ^[26, 27]^.

In conclusion, by comparative analysis of genome-wide gene expression data from the public domain, we show, firstly, that the common transcriptional response of *E. coli* to stress is represented by down-regulation of energy-expensive flagellar assembly pathway and reciprocal up-regulation of the global regulator rpoS. Secondly, that biocides and antibiotics are associated with distinct gene expression programs. Thirdly, that excess copper, at near lethal concentration, behaves as an antibiotic at the level of transcriptomic response elicited in *E. coli*. Copper may be used synergistically along with other antibacterials for better results as revealed by other studies ^[28, 29]^. Lastly, the reciprocal relationship between spermidine and putrescine uptake genes is a novel and interesting finding which warrants further study.

## Acknowledgements

RC was supported by a grant from Department of Science and Technology, New Delhi, India (grant no: SR/WOS-A/208/2009). SKM thanks Dr. Debasisa Mohanty, National Institute of Immunology, New Delhi, India for critical suggestions during data analysis.

---

**Figure S1.**
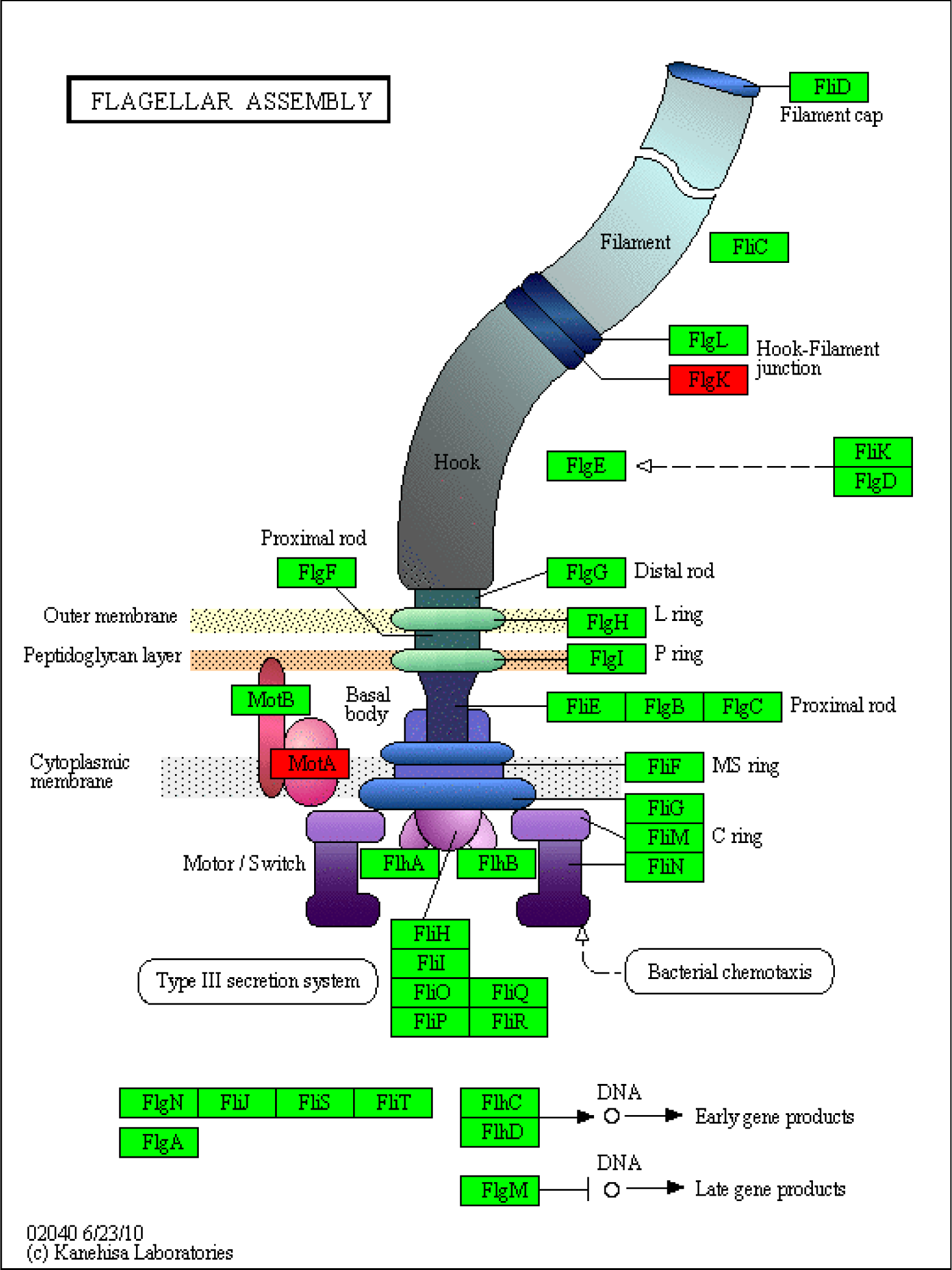
This figure was generated by feeding the list of NCBI GeneIDs belonging to the pathway “Flagellar assembly” at http://www.genome.jp/kegg/tool/color_pathway.html, searching against the genome *E. coli* K12 MG1655. Green color of the node suggests negative relative expression of that gene, i.e., down-regulation, red color suggests the opposite. The figure shows that most genes participating in Flagellar biosynthesis are down-regulated under stress.

